# Committed Hemopoietic Progenitors, Not Stem Cells, are the Principal Responders to Hox Gene Transduction

**DOI:** 10.1101/174490

**Authors:** Harvey Lim, Salima Janmohamed, Patricia Benveniste, Robert Herrington, Mary Barbara, Catherine Frelin, Deborah Hyam, Christopher J. Paige, Juan-Carlos Zuñiga-Pflücker, Carol Stocking, Jana Krosl, Guy Sauvageau, Norman N Iscove

## Abstract

As hemopoietic stem cells differentiate, their proliferative lifespan shortens by unknown mechanisms. Homeobox cluster (Hox) genes have been implicated by their enhancement of self-renewal when transduced into hemopoietic cells, but gene deletions have been inconclusive because of functional redundancy. Here we enforced HOXB4 expression in purified precursor stages, and compared responses of early stages expressing the endogenous genes with later stages that did not. Contrary to the prevalent view that transduced Hox genes enhance the self-renewal of hemopoietic stem cells, stem cells or their multipotent progeny expressing the endogenous genes showed little response. Instead, immortalization, extensive self-renewal and acquired reconstituting potential occurred in committed erythroid and myeloid progenitors where the endogenous genes were shutting down. The results change our understanding of the stages affected by exogenous HOX proteins and point to shutdown of the endogenous genes as a principal determinant of the shortened clonal lifespans of committed progenitor cells.

## INTRODUCTION

Proliferative hierarchy is a signature feature of the hematopoietic system. Stem cells having indefinite regenerative and proliferative capacity differentiate through downstream stages characterized by progressively shortening clonal lifespans. Proliferative lifespan is linked to the concept of "self-renewal", a defining capacity of stem cells to generate daughter cells that retain the indefinite regenerative capacity of the parent as well as other daughters that progress into lineage divergence, functional maturation and cessation of growth. Despite much descriptive study, and identification of numerous individual genes associated with gain or reduction of precursor cell function in overexpression or deletion analyses, no coherent understanding has yet emerged to explain how various players are coordinated to specify or govern proliferative lifespan.

The classical homeobox cluster (*Hox*) genes, known to govern segmental and cellular identity in embryonic development, originally attracted attention as potential regulators of the stem cell state when they were found to be expressed in primitive hemopoietic cells but not in terminally maturing progeny (Pineault et al., 2002; Argiropoulos and Humphries, 2007). The *Hox* gene family is represented in mammals by 4 separate but analogously organized clusters encoding a set of 39 paralogous proteins in total, each possessing an essentially invariant homeodomain (Shashikant et al., 1991; Mark et al., 1997). Various paralogs have been tested for their potential role in hemopoietic precursor cells by examining the effect of transduction and enforced expression of individual paralogs in hemopoietic precursor cells (Sauvageau G et al., 1995). Most were found to promote dramatic increase in precursor cell self-renewal and marrow regenerative capacity in transplantation assays (Sauvageau G et al., 1995; Sauvageau et al., 1997; Thorsteinsdottir et al., 1997, 2002; Fischbach et al., 2005; Schiedlmeier et al., 2007; Iacovino et al., 2009; Bach et al., 2010; Fournier et al., 2012). All of these reports attributed the observations to enhanced self-renewal occuring in multipotent stem cells.

While these earlier results hinted at a possible involvement of the *Hox* family in the process of self-renewal, they were inconclusive. The observed effects could have reflected anomolous responses to supraphysiological amounts of HOX protein, sequestration of cofactor elements, or imbalanced expression relative to other *Hox* elements. For confident attribution of function and requirement of the endogenously expressed genes, it has therefore been critical to determine the consequences of complete withdrawal of the endogenous HOX proteins. A number of studies have examined inactivation of individual *Hox* genes. Not surprisingly given the extensive functional redundancy in this system, inactivation of individual elements generally yielded ambiguous phenotypes with substantial residual stem cell function, and even haploid deletion of the entire A or B clusters had surprisingly mild effect on sustained hematopoiesis (Lawrence et al., 1997; Björnsson et al., 2003; Brun AC et al., 2004; Bijl J et al., 2006; Magnusson et al., 2007; Lebert-Ghali et al., 2010).

Here we describe experiments designed to shed new light on the potential role of the endogenous Hox cluster genes by defining more clearly the relationship between expression of the endogenous Hox genes in various precursor cell stages and the responsiveness at each stage to enforced expression of an exogenous Hox transgene.

## RESULTS

### HOX genes are coordinately expressed through early stem cell stages until shutdown in committing erythroid and myeloid progenitors

Expression of A and B cluster Hox genes in both human and mouse HSC has been thoroughly documented. However, the extent to which expression persists in differentiating progeny has not been clearly defined. Pineault et al. (Pineault et al., 2002) showed expression of a panel of A and B cluster transcripts in the Sca1+Lin-fraction of murine marrow enriched in primitive precursors including HSC, but only low to undetected levels in fractions expressing lineage markers of maturation. Using a Hoxb4-YFP reporter expressed from the native promoter, Hills et al. (Hills D et al., 2011) found most cells in the Kit+Sca1+Lin-fraction including functionally assayed HSC to express the fusion protein. Since this fraction also contains more advanced progenitor stages, the finding suggested that Hoxb4 expression may continue in differentiating HSC progeny. However, precursor stages further downstream of the Kit+Sca1+Lin-fraction were not analyzed in detail. Thus, between the HSC and primitive progenitor cells which express the Hox cluster genes and the advanced lineage marker-positive cells which no longer express them, the available studies have left unresolved the particular stages at which the Hox genes are silenced. Since that information would likely contribute to an understanding of the role played by the endogenous Hox genes, we elected to map their expression over a detailed series of stages from the earliest stem cells through lineage-committed progenitors to terminally differentiated blood cells.

The analysis resolved the following stages in murine bone marrow. The most primitive cells are the long-term multipotent stem cells LT-HSC. These sustain self-renewal and thus systemic blood cell production through serial transplantations over timespans that exceed the lifetime of a single mouse (Iscove and Nawa, 1997). LT-HSC are purified to absolute functional homogeneity by cell sorting on the basis of low Rhodamine123 dye retention, expression of surface c-Kit, Sca-1 and SLAMF1/CD150, and minimal expression of α2 integrin/CD49b and a suite of mature hematopoietic lineage markers (Frelin et al., 2013), a phenotype summarized here using the nomenclature RKSL_CD34^-^_SLAMF1^+^_α2^lo^. Phenotypically distinct Intermediate-Term IT-HSC (RKSL_CD34^-^_SLAMF1^+^_α2^hi^, differing in expression of α2 integrin) are, like LT-HSC, capable at single cell level of regenerating all hematopoietic lineages in a mouse but do not sustain self-renewal or multilineage blood cell production beyond 4 months (Benveniste et al., 2010). Single IT-HSC are estimated to generate 10^12^ cells over their proliferative lifetime.

Downstream of the IT-HSC are further differentiated progeny which have further diminished proliferative lifespans and in which lineage restriction begins. Short-term ST-HSC are isolated on the basis of upregulated CD34 expression and absence of the Flt3 receptor that marks early lymphoid precursors (KSL_CD34^+^_Flt3^-^) (Yang et al., 2005). They sustain blood cell production for only 4 wk, generating a lifetime clonal output of only 10^8^ cells. Beyond the primitive Kit+Sca1+Lin-population, a more advanced mixed "Progenitor" fraction marked by expression of c-Kit and CD34 but not Sca-1 (KS^-^L_CD34^+^_Flt3^-^)(Akashi et al., 2000; Terskikh et al., 2003) has even more fleeting regenerative potential and a clonal output capped at about 10^5^ - 10^6^ cells. Transcript expression in more advanced stages was measured in a series of cDNA samples derived from partially and completely committed erythroid and myeloid precursors whose identity and homogeneity were validated functionally in colony assays (Brady et al., 1995; Billia et al., 2001).

LT-and IT-HSC are in a quiescent state in normal marrow, whereas ST-HSC and more advanced progenitors actively cycle (Wilson et al., 2008; Foudi et al., 2009; Oguro et al., 2013). To detect possible linkage of gene expression with cycling status, we analyzed not only quiescent, freshly isolated LT-and IT-HSC ("Fresh"), but also the same purified cells after they were induced into active cell cycle in culture ("Cultured"), at time points (LT-HSC 48 hr; IT-HSC 36 hr) at which about 50% of cells had divided (Benveniste et al., 2010). Transcript expression was quantitated for a selection of *Hox* genes sampled over the 5’ to 3’ extent of the *Hox* A and B clusters. Overall, shown in **Figure 1a**, cluster members at positions 3, 4, 6, 9 and 10 were expressed in both quiescent and cycling LT-and IT-HSC, with exceptions showing quantitative cycle-associated differences highlighted in **Figure 1b**. Expression continued at comparable levels through ST-HSC. Somewhat surprisingly, this level of expression was sustained even in Progenitor fraction cells (K+S-L-). Thereafter, *Hox* 3 and 4 transcripts abruptly diminished in cells committing into myeloid and erythroid branches of development. Expression of *Hox* A cluster 5, 9 and 10 transcripts however was maintained even in erythroid/megakaryocyte (BFUE, *Hox a5* and *a10*), megakaryocyte (*a9*), neutrophil/macrophage (*a5,9,10*) or pure macrophage (*a5,9*) colony-forming cells. Finally, expression was no longer detected in more advanced pure neutrophil or erythroid colony-forming cells. Expression of at least *Hox4* paralogs also persisted in B and T lymphoid precursor cells (**Figure 1**), known for their extensive proliferative capacity in primary immune responses and establishment of ’memory’ reservoirs. Thus, expression of transcripts for the tested *Hox* cluster paralogs was sustained well beyond the K+S-L-mixed progenitor fraction, and ceased only in advanced precursor cells restricted to individual erythroid or myeloid lineages.

**Figure 1.**
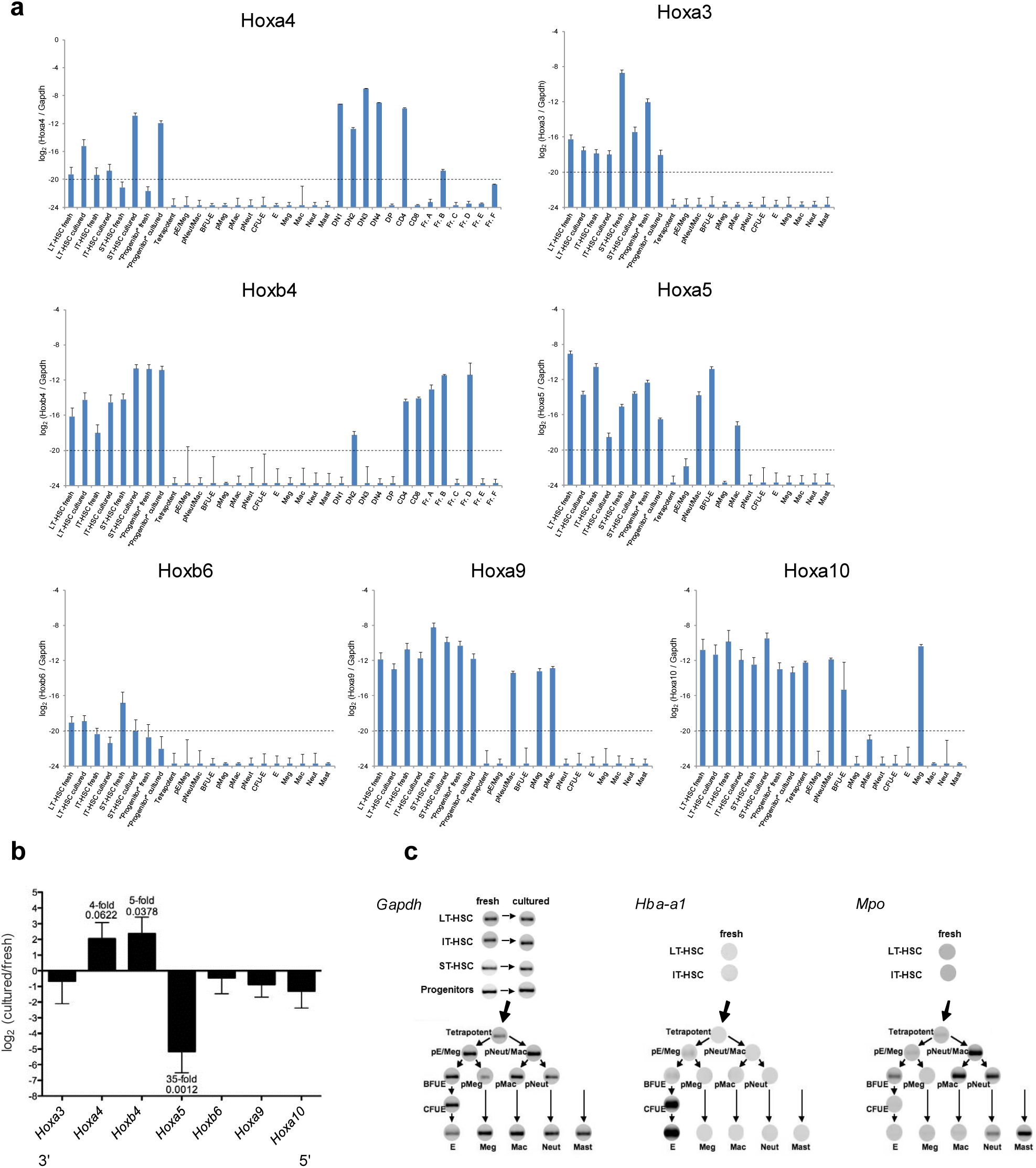
Transcript-level expression of selected Hox genes in the hematopoietic precursor cell hierarchy. **(a)** Quantitative PCRs were performed on globally amplified cDNAs generated from each stratum of the hematopoietic hierarchy. cDNA samples at each precursor stage were pooled from 6 - 12 biological repeats (purification and preparation of cDNA). Means and SEMs of triplicate Q-PCR determinations on each pool are shown. Dashed lines represent the limit of detection. Precursor nomenclature: Tetrapotent: Erythroid/Megakaryocyte/Neutrophil/Macrophage (E/Mac/Neut/Meg) precursors; p"lineage": bipotent and unipotent precursors of the designated lineages. E, Meg, Mac, Neut, Mast cDNAs were sampled from mature cells in pure lineage clones grown in culture. **(b)** Cycle-related expression of selected *Hox* transcripts in HSC. Q-PCR results were pooled from 12 and 10 independent experiments for fresh and cycling LT-and IT-HSC respectively. Log_2_ ratios of the mean expression of cultured samples to the mean expression of fresh samples are plotted with standard errors. p values were calculated using log_2_-transformed data by an unpaired 2-tailed t test. The fold difference and p value is shown for transcripts differing strongly between resting and cycling states (where p<0.05 or p<0.10 and fold difference > 2^2^). **(c)** Semi-quantitative PCR bands from agarose gels are shown for *gapdh* and lineage specificity control (*Hba-a1*, *Mpo*) transcripts at their respective positions in the hierarchy. PCR reactions for each gene were cycled until bands of the anticipated size were visible on agarose gels, between 30 and 45 cycles.

### Enforced expression of a *Hox* transgene renders IT-HSC capable of permanent multilineage reconstitution

The correlation of *Hox* gene expression with proliferative potential led us to consider that the shortened proliferative lifespan of cells downstream of the long-lived LT-HSC might be connected with the eventual withdrawal of the HOX cluster proteins. A critical test would be to ask whether normally transient downstream progenitors would sustain permanent in vivo reconstitution if the availability of HOX protein could be artificially sustained. We performed our initial tests on purified IT-HSC, which normally sustain blood cell production in all lineages for 12-16 wk before the clones regress. Single freshly purified cells were infected in culture with a retrovirus enforcing expression of *HOXB4*, a representative *Hox* paralog that enhances self-renewal as a transgene with minimal effect on output of differentiated blood cells or primary leukemogenic potential. After infection and further culture for a total of 8 d, transduced clones were transplanted into gamma-irradiated recipient mice and engraftment with donor blood cells was followed for 32 wk (**Figure 2a**). Of 37 single IT-HSC exposed to *HOXB4* retrovirus, 14 regenerated erythropoiesis in recipient mice to high level donor composition and sustained red cell production through 32 wk. In contrast, no reconstituting activity survived the culture period when cells were infected with control retrovirus lacking a *HOXB4* insert, indicating that the culture conditions were adverse to maintenance of normal functional IT-HSC.

**Figure 2.**
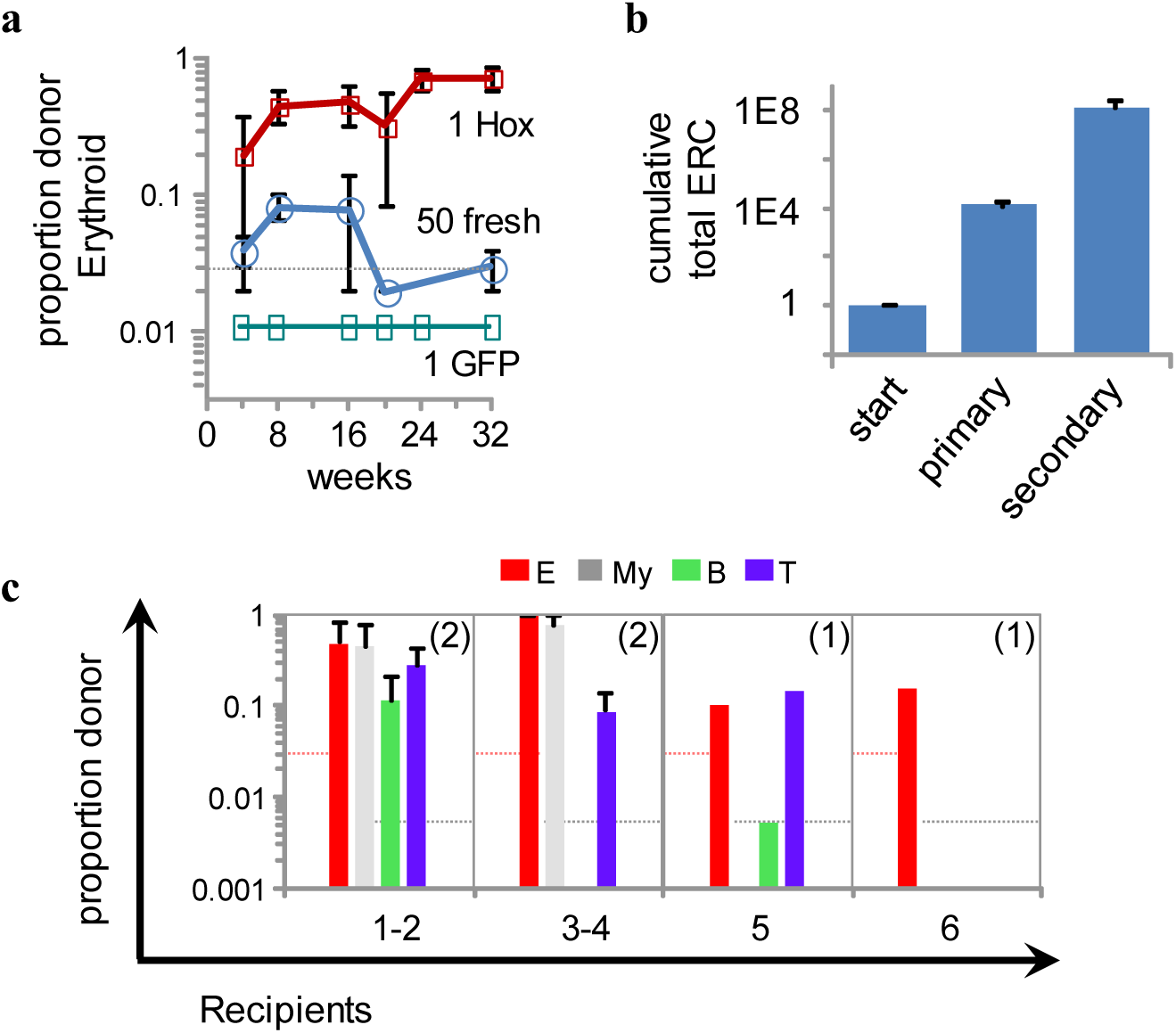
Acquisition of long-term reconstituting potential by IT-HSC following *HOX* gene transduction. All reconstitutions are plotted as proportion (not %) of the designated lineage contributed by the engrafted cells, in this and subsequent figures. **(a)** Single IT-HSC were transduced in four independent experiments and each of 37 resulting clones was injected into a lightly irradiated *Kit*^*W-41*^ recipient along with 5x10^5^ recipient Gpi1-type *Kit*^*W-41*^ marrow cells. Shown are the averaged kinetics of erythroid reconstitution in 14 successfully engrafted mice, from single HOX-transduced cells ("1 Hox"), or from 50 freshly isolated IT-HSCs within the same experiments, or from single control virus-transduced cells ("1 GFP"). **(b)** Marrow was harvested from the primary recipients of *HOX*-transduced IT-HSC shown in (a) at 33 weeks post-transplant, and 10^4^ cells were injected into lightly irradiated *KitW-41* recipients. The cumulative numbers of Erythroid Reconstituting Cells (ERC) are plotted. Primary mice used for secondary passage contained grafts composed of at least 90% donor erythroid cells. 30 wk after transplant, marrows from four secondary recipients were transferred into tertiary recipients. The proportion of donor erythrocytes in positive secondary recipients averaged 54%. Recipients were scored for presence of donor erythrocytes at 24 weeks post-transplant. **(c)** Lineages reconstituted in 6 primary recipients of single *HOX*-transduced IT-HSC at 22-30 weeks post-transplant. The bars indicate mean proportion chimerism in each lineage, with SEMs showing variation between single recipients of individual clones and numbers of recipients indicated in parentheses. The dashed lines represent detection thresholds for donor erythroid proportions quantitated in Gpi1 assays, and for the other lineages measured by flow cytometry.

To test further for durability of the original erythroid clones, marrow from primary recipient mice was passaged into irradiated secondary recipients. Passage at limiting cell numbers -12 of 26 recipients of 10000 cells regenerated erythrocytes of donor origin -allowed estimate of the frequency of cells with erythroid reconstituting capacity in the primary marrows at around 1 in 16000 cells, and of their absolute number per mouse (assuming 2 x 10^8^ total marrow cells) at about 12000 representing a 12000-fold expansion from a single injected reconstituting cell. After 30 wk, marrow cells from secondary recipients were similarly passaged into tertiary recipients, revealing a similar frequency of erythroid reconstituting cells in the secondary marrows and a further 10000 fold-expansion in number of reconstituting cells. The cumulative expansion of reconstituting cell precursors over the course of 64 wk was about 120 million-fold starting from a single originally transduced IT-HSC (**Figure 2b**), and red cell production continued unabated in tertiary recipients for an additional 32 wk. The differentiation potential of single transduced IT-HSCs into all hematopoietic lineages was tracked in 6 mice. While each of the erythroid, myeloid, B and T cell lineages regenerated from donor cells in 2 of the mice, 4 other recipients sustained production in only a subset of lineages 4 -6 months post-transplant (**Figure 2c**).

The result showed that enforcement of the availability of HOX protein endowed clones deriving from normally transient single IT-HSC with the capacity to sustain blood cell production over at least a mouse lifetime, accompanied by dramatic expansion in numbers of cells able again to reconstitute red cell production in mice at single cell level. Further, the expansion of erythroid reconstituting cell numbers occurred in concert with production of mature erythrocytes and without expansion of reconstituting cell frequencies, implying continued responsiveness of transduced progenitors to external regulation.

### Permanent multilineage reconstitution from transduced ST-HSC

In contrast to IT-HSC which can regenerate hemopoiesis in recipient mice from a single injected cell, 100’s of freshly isolated ST-HSC must normally be injected for detection of multilineage systemic grafts, which peak at 3 wk and then regress (**Figure 3a**). ST-HSC, in groups of 10 or singly, were retrovirally transduced and injected into irradiated mice. In contrast to the performance of freshly isolated and untreated cells, even single transduced cells yielded high level reconstitutions in 7 of 23 recipient mice. Grafts were sustained at least 20 wk in individual mice in erythroid, myeloid, and B and T lymphoid lineages (**Figure 3a**). Although all lineages were represented collectively in the engrafted recipients of single transduced cells, only a subset of lineages was represented in most individual recipients (**Figure 3b**), recapitulating the results already seen with individual transduced IT-HSC. The results strengthened the evidence that enforced availability of HOX protein could confer sustained reconstituting capacity upon more advanced cells that had already begun the process of lineage restriction. They also showed that persistence of HOX expression did not preclude differentiation through any of the erythroid, myeloid, B or T lymphocye lineages.

**Figure 3.**
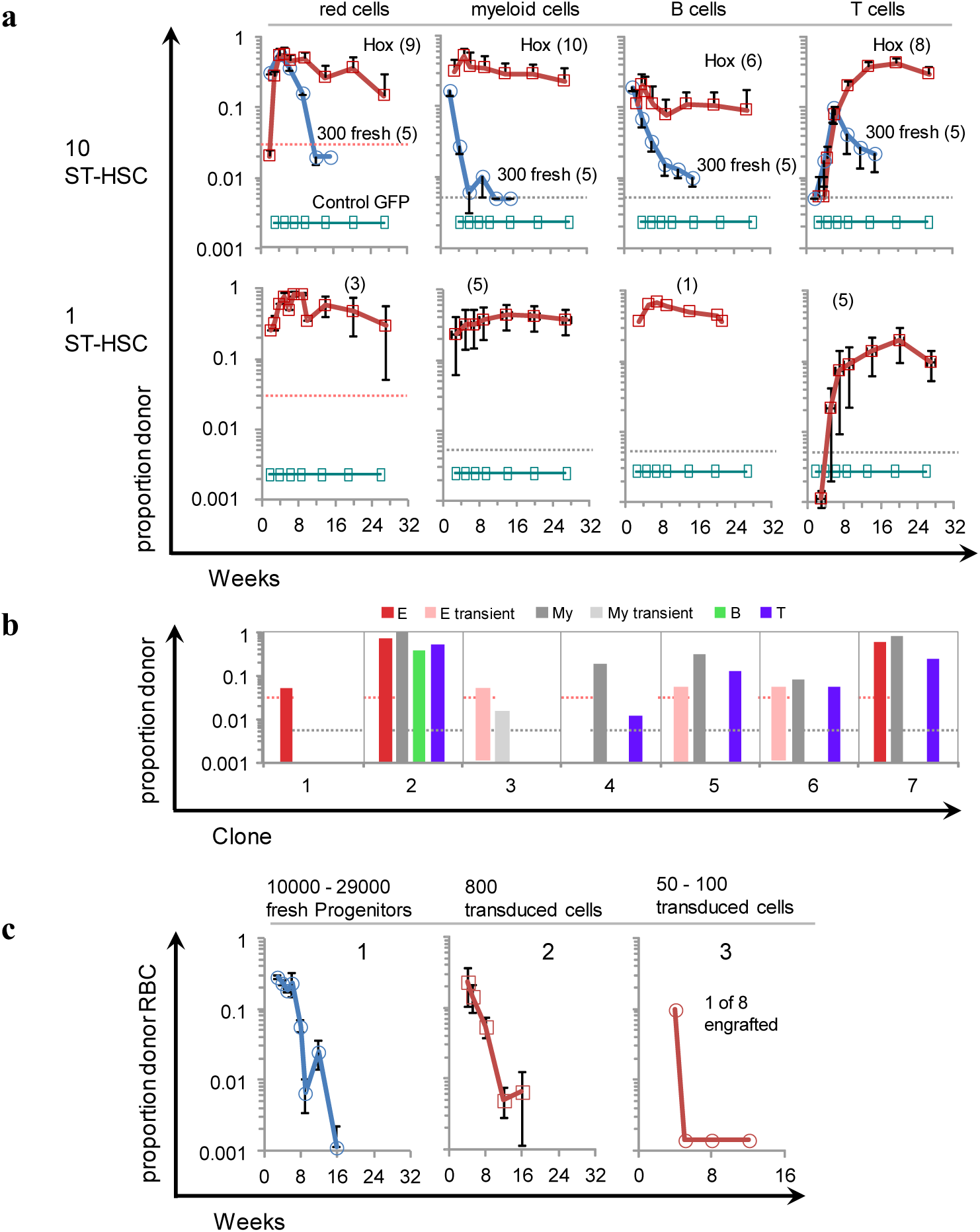
Enforced expression of a *HOX* gene endows ST-HSCs but not Progenitors with long-term reconstituting capacity. **(a)** Freshly sorted ST-HSC, 300, or cultures derived from 10 or single ST-HSC infected with control (GFP only) or *HOX*-expressing retrovirus were each transplanted into a lethally irradiated C57B6/J mouse along with 1 x 10^6^ *Kit*^*W-41*^ host-type Gpi1 marrow cells. Cultures with 10 cells were initiated in 3 independent experiments and totals of 5 control-infected and 11 *HOX*-transduced cultures were tested in host mice. Single cell cultures were initiated in 2 independent experiments and totals of 10 control-infected and 23 *HOX*-transduced cultures were tested. The average proportion of donor cells in blood red cell, myeloid, B and T lymphocyte lineages is shown with SEM. Averages for each lineage represent only those mice (numbers shown in parentheses) that contained a detectable graft in the lineage plotted. Dashed lines show detection thresholds. **(b)** Donor representation in blood cell lineages in 7 individual recipients of single transduced ST-HSC after 20-24 wk. Lineages present transiently at earlier time points are also indicated. **(c)** Donor erythroid cells in recipients of *HOX*-transduced Progenitor fraction cells. Panel 1, mean and SEM of proportion donor erythrocytes after transplant of 10000 - 29000 transduced cells into 12 mice in 4 independent experiments. Panel 2, mean and SEM of proportion donor erythrocytes in 9 recipients of 800 transduced cells in 4 independent experiments. Panel 3, proportion donor erythrocytes in 1 of 8 recipients of 100 transduced cells in a single experiment; blood myeloid, B and T lineage cells of donor origin were not detected. 7 other recipients had no detectable graft.

Progenitor fraction cells lack the proliferative capacity of ST-HSC, and 1000’s of freshly isolated control cells need to be injected in order to detect low levels of systemic engraftment in a 2 – 4 wk interval. When HOX expression was enforced by retroviral transduction into purified Progenitor fraction cells, a small subset of the cells acquired an enhanced capacity for initial and fleeting engraftment, but reconstitution was not sustained (**Figure 3c**) even though cultures were positive for the retroviral GFP marker at the time of injection into recipient mice.

### Multilineage reconstitutions from multipotent precursors are dominated by transplantable progenitors that are lineage restricted

Enhanced multilineage reconstitutions from *HOX* gene-transduced marrow precursors is currently understood to reflect increased regenerative and self-renewal activity of multipotent HSC, manifesting as net expansion of their numbers in regenerating marrow of recipient mice. We therefore expected to be able to identify large numbers of functional multipotent HSC in the bone marrows of mice that had been transplanted with *HOX*-transduced multipotent cells. As expected, mice transplanted with 30 *HOX*-transduced LT-HSC regenerated in erythroid, myeloid, B and T cell lineages, and passage of 2 x 10^6^ of their marrow cells into secondary recipients similarly yielded grafts containing all lineages. As shown in **Figure 4a,** 5 primary recipients of single transduced LT-or ST-HSC also reconstituted in all lineages. Three of these recipient marrows were analyzed by passage at limiting dilution into secondary recipients in 3 independent experiments. Of a total of 96 secondary recipients, 28 engrafted and 27 of the grafts consisted of erythroid/myeloid or myeloid cells only. Limiting dilution clones containing lymphoid cells were rare. In 120500 total cells sampled, including additional experiments not shown in the figure, T lymphocytes were present in 3 recipients, while B lymphocytes were only seen in 3 of 6 recipients of 2 x 10^6^ cells. Collectively, the observations imply the presence in *HOX*-transduced primary multilineage marrows of erythroid and/or myeloid reconstituting cells at frequencies ranging from 1:2000 to 1:20000 marrow cells, of reconstituting cells with T lymphoid potential near 1:40,000, and of repopulating cells with B lymphoid potential at a frequency closer to 1:2,000,000. These estimates imply that the frequency of individual multipotent *HOX*-transduced cells able to reconstitute all lineages would not have been higher than 1:2,000,000, well below the 1:20000 predicted if there had been extensive increase in numbers of multipotent HSC during marrow regeneration up to their frequency in normal marrow.

**Figure 4.**
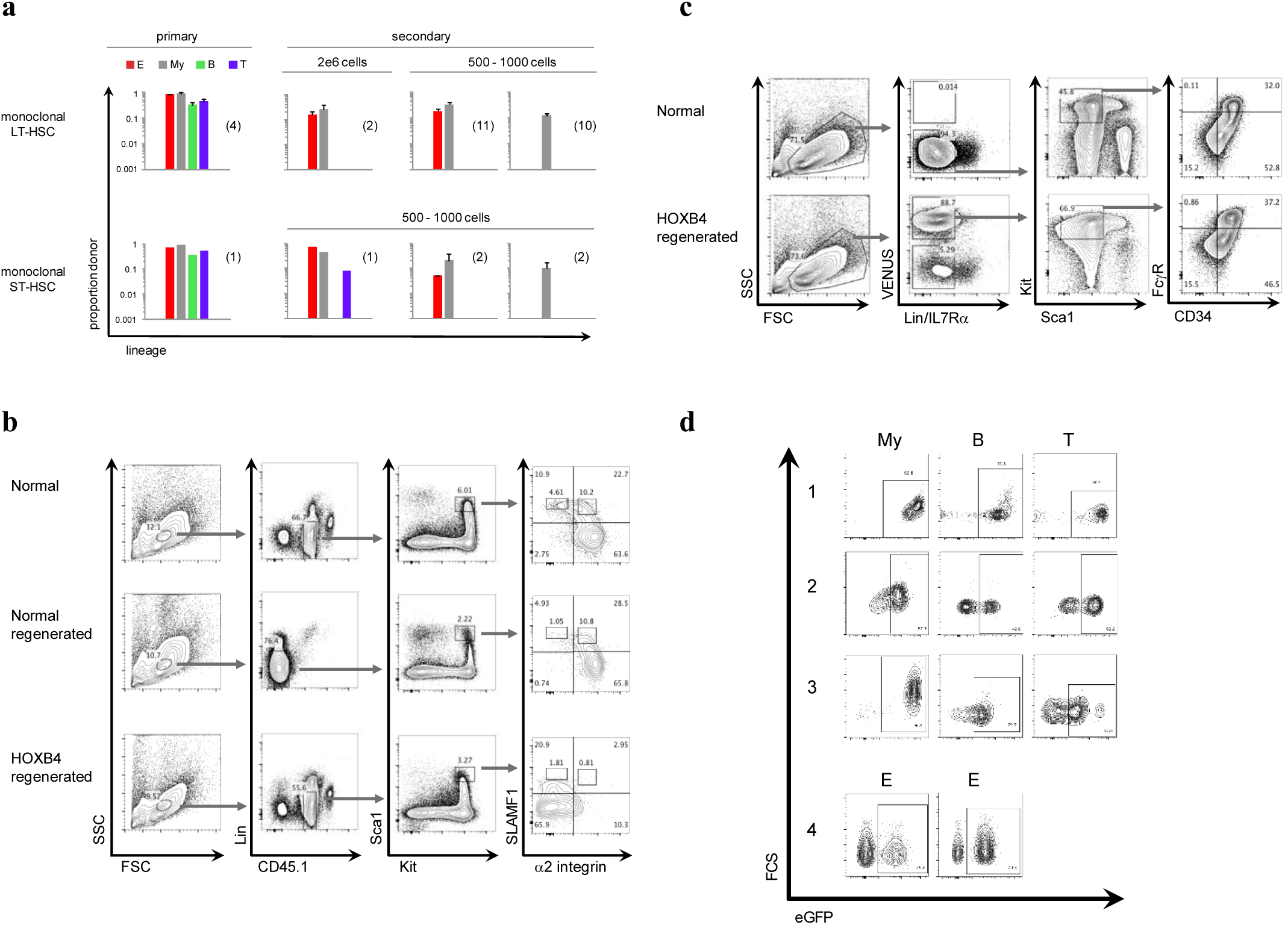
Functional capacities and phenotype of cells present in marrow of multilineage reconstituted recipients of *HOX*-transduced cells. Donor representation in all blood cell lineages of primary recipients 20 wk after transplantation of transduced LT-and ST-HSC, and lineage-restricted 20 wk reconstitutions in secondary recipients of limiting numbers of marrow cells from the multilineage-reconstituted primary recipients. Average proportions of donor contribution are indicated with SEM for each lineage. Numbers of individual mice are shown in parentheses. "Monoclonal" primary recipients were reconstituted from a single *HOX*-transduced cell. **(b)** Flow cytometry profiles, representative of 3 independent analyses, of normal CD45.1 marrow, marrow from host mice 20 wk after transplant with 1 x 10^6^ normal CD45.2 marrow cells ("normal regenerated"), or marrow from mice 10 - 20 wk after transplant with 30 *HOX*-transduced CD45.1 LT-HSC. Arrows show the originating gate for each profile. The gates containing highly purified functional LT-HSC (Lin^-^_cKit^+^_Sca1^+^_SLAMF1^+^_α2^lo^) and IT-HSC (Lin^-^_cKit^+^_Sca1^+^_SLAMF1^+^_α2^hi^) in normal marrow are indicated in each SLAMF1/α2 integrin profile. **(c)** Flow cytometry profiles representative of 2 independent analyses of normal marrow or marrow from mice 44 - 49 wk after transplant of a single *HOX*-transduced IT-HSC clone. Transduced cells coexpress VENUS fluorescent protein. Myeloid progenitors (Kit^+^Sca1^-^Lin^-^IL7Rα^-^CD34^hi^FcγR^hi^) represented 0.16% versus 0.55% of normal versus *HOX*-transduced marrow respectively, and erythroid progenitors (Kit^+^Sca1^-^Lin^-^IL7Rα^-^CD34^lo^FcγR^lo^) 0.08% versus 0.23%. **(d)** Flow cytometry profiles showing expression of retroviral fluorescent marker in blood cells of donor Ly5 type in the indicated lineages. **1**, recipient of transduced cells originating from 30 LT-HSC, at 23 wk; **2**, secondary recipient at 26 wk of marrow from the mouse shown in 1, or 49 wk in total after transduction; **3**, recipient of transduced cells originating from a single LT-HSC, 29 wk; **4**, red cell fluorescence in recipients of transduced cells. Left, primary recipient of 5 x 10^5^ transduced marrow cells at 26 wk; Right, secondary recipient at 13 wk of 2 x 106 marrow cells from a similar primary recipient, or 39 wk after the original transduction.

### Absence of phenotypic LT-or IT-HSC and elevated progenitor numbers in recipients of transduced LT-HSC

Given the rarity of fully multipotent cells in marrow of mice regenerated in all lineages from transduced HSC, it was of considerable interest to ask whether we could identify cells having the phenotype of HSC by flow cytometry. **Figure 4b** compares phenotypes of marrow from mice that were originally transplanted with either untransduced normal marrow or with 30 transduced LT-HSC. The LT-HSC and IT-HSC gates in the right-most column contain highly purified functional HSCs in normal marrow, and the same windows were also populated in marrow regenerated from normal marrow cells. In contrast, in marrow regenerated in all lineages from transduced LT-HSC, these gates were practically empty, containing events only at their lower periphery, and enduring grafts were not obtained from cells in either of the entire Kit^+^_Sca1^+^_Lin^-^_α2 integrin^-^_SLAMF1^+^ or SLAMF1^-^ fractions (not shown). The absence of phenotypic HSC in numbers comparable to those in mice regenerated from normal marrow again suggested that no detectable increase in multipotent HSC numbers occurred during marrow regeneration as a consequence of *HOX* gene transduction. The outcome reinforced the result of the functional analysis which similarly demonstrated that multipotent cells in *Hox*-regenerated marrow were rare if present at all. Despite the apparent absence of phenotypic HSC, phenotypic myeloid (Kit^+^Sca1^-^Lin^-^IL7Rα^-^CD34^hi^FcγR^hi^) and erythroid (Kit^+^Sca1^-^Lin^-^IL7Rα^-^CD34^lo^FcγR^lo^) progenitors of donor origin in recipients of *HOX*-tranduced IT-HSC clones were present in 2 - 3-fold greater numbers than in normal control marrow (**Figure 4c)**.

Flow cytometry provided additional information on the stability and uniformity of expression of the retroviral cargo. **Figure 4c** shows that the elevated numbers of erythroid and myeloid progenitors expressed high levels of the fluorescent retroviral expression marker. Expression of the fluorescent marker in at least half of the mature blood cell progeny of transduced precursor cells in each of the erythroid, myeloid, B and T cell lineages is shown in **Figure 4d**. Expression in mature cells provides evidence for relatively unbiased expression of retroviral cargo in precursor cells in all these lineages. Proportions of red cells that were fluorescent corresponded closely with proportions of donor red cells measured by GPI1 isotype, implying persistence of retroviral expression in all transduced red cell precursors.

### Dependence of *HOX*-immortalized clones on continued availability of the *HOX* transgene

To evaluate further the relationship between HOX expression and proliferative lifespan, we asked whether the extended lifespan of the *HOXB4*-transduced grafts depended on the continued supply of exogenous HOX protein. Exogenous *HOXB4* could simply exert the same self-renewal-promoting role as the endogenously expressed counterparts, and its enforced expression would simply ensure the continued availability of HOX protein. In that case, grafts from transduced precursor cells should be addicted to HOXB4 and should fail to sustain long term reconstitution if the exogenous transgene were removed. Alternatively, exogenous HOXB4 protein, perhaps in supraphysiological amounts, could irreversibly "reprogram" progenitor cells in analogy to the iPS paradigm and in a way that stabilizes the self-renewing state. In that case, the grafts should continue to self-renew and regress after removal of the transgene.

To distinguish between these alternatives, we constructed a HOXB4 retrovirus which would permit Cre-mediated deletion of its cargo when desired by virtue of LoxP sites located in the flanking proviral LTRs (**Figure 5a**). Cells from ER-Cre mice release cytoplasmically sequestered CRE recombinase to the nucleus on exposure to tamoxifen (Hayashi and McMahon, 2002). We infected LT-or IT-HSC purified from ER-Cre marrow with HOXB4-VENUS-LoxP retrovirus and transplanted them into irradiated mice. After polyclonal regeneration of marrow by the infected cells, recipient mice were treated with tamoxifen (**Figure 5b**). As shown in **Figure 5c**, VENUS fluorescence was substantially reduced in circulating blood cells in all hosts, accompanied by early loss of donor myeloid and erythroid components and more gradual loss of donor B and T lymphocytes. Tamoxifen-treated grafts were unable to engraft secondary marrow recipients. In contrast, grafts of cells infected with Hoxb4-VENUS virus lacking LoxP sustained high level donor reconstitution despite tamoxifen treatment and robustly reconstituted secondary recipients. These results show that persistence of reconstituting clones in primary recipients and their capacity to reconstitute secondary recipients remained dependent on the continued expression of the *HOXB4* transgene. As shown in **Figure 4a-c**, the self-renewing elements in primary grafts are precursors possessing the phenotype and characteristic lineage restriction of advanced progenitor cells in which endogenous Hox cluster genes cease to be expressed.. The reversibility of the immortalizing effect therefore shows that it is the exogenous HOXB4 that maintains the self-renewing capacity of these cells, and that irreversible reprogramming does not take place.

**Figure 5.**
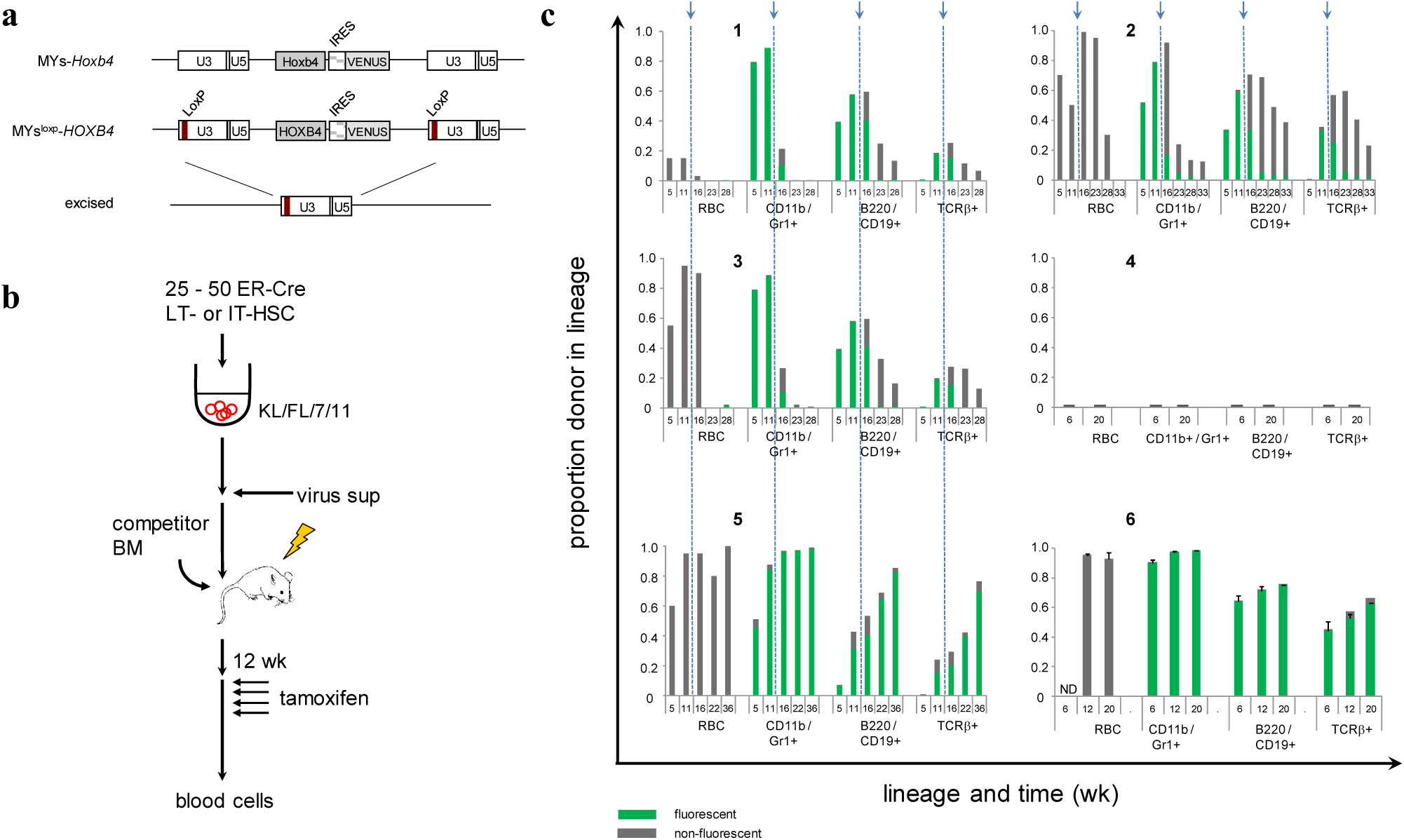
Effect of withdrawal of exogenous Hoxb4 on persistence of hemopoietic reconstitution. **(a)** Structure of the MYs^loxp^-*HOXB4* retrovirus showing the integrated provirus with LoxP sites in both 5’ and 3’ LTRs, and the control MYs-*Hoxb4* construct lacking LoxP sites. Cre recombinase deletes the proviral contents as shown, leaving behind a single copy of the retroviral LTR. **(b)** After culture of 25 - 50 purified ER-Cre LT-or IT-HSCs for 30 hr, supernatant medium from cultured virus producer cells was added. Wells containing cells visually confirmed to be VENUS^+^ were harvested on day 8 and transplanted polyclonally into irradiated mice with kit^W41/W41^ marrow competitor cells. At 12 wk recipient mice received tamoxifen on 5 successive days for release of CRE recombinase in the transplanted ER-Cre cells. **(c)** Proportion of donor cells in individual primary recipient mice in the indicated lineages at the indicated time points. Arrows and dashed lines indicate the time point of tamoxifen treatment. Panels 1, 2 and 3 are representative of 14 recipient mice reconstituted with MYs^loxp^-*HOXB4* infected cells, showing high levels of initial reconstitution by virus-infected cells, depletion of VENUS fluorescence in all lineages after tamoxifen treatment, steep loss of donor erythroid and myeloid and slower fall-off of B and T lymphoid graft components. Panel 4 shows the result of transplant of 10^7^ marrow cells at 28 wk from the tamoxifen-treated recipient in panel 3 into 2 secondary recipient mice, indicating the absence of detectable engraftment in all lineages. Panel 5 shows reconstitution in all lineages from cells infected with MYs-*Hoxb4* virus lacking loxP and absence of effect of tamoxifen treatment (representative result of 15 recipients). The results in panels 4 and 5 were obtained in 3 independent experiments. Panel 6, vigorous reconstitution in all lineages after passage of 24 wk marrow from the primary mouse shown in Panel 5 into secondary recipients. ND = no data. Aggregate results are representative of 4 independent experiments.

## DISCUSSION

This analysis was undertaken to advance an understanding of the role of the endogenous homeobox cluster genes in controlling self-renewal and proliferative lifespan in the hemopoietic stem and progenitor cell hierarchy. Our initial intent was to test for a possible link between shutdown of expression of the homeobox genes and the characteristic truncated clonal lifespans of grafts initiated by IT-and ST-HSC in irradiated recipient mice. Our experiments showed that enforced expression of a HOXB4 transgene in these precursors indeed rendered them capable of initiating multilineage grafts that endured long-term. The observations were consistent with the notion that IT-and ST-HSC had themselves been affected by the transgene, responding with enhanced and sustained self-renewal. However, subsequent findings turned out not to support the initial interpretation.

Two new and critical insights emerged from the rest of the analysis. First, high resolution mapping of a selection of Hox cluster genes documented their RNA-level expression from multipotent HSC down to and including mixed Sca1^-^ progenitor cells. Thereafter, Hox 3 and 4 paralogs shut down at BFU-E/Meg and GM-CFC stages, followed by extinction of Hox 5, 9 and 10 in yet more advanced monopotent precursors. These results raised the question of whether a transduced homeobox gene could influence early cell stages in which multiple endogenous homeobox genes were already actively expressed. Second, marrows that were reconstituted in all lineages from HOXB4-transduced precursor cells were analyzed by both biological assay at limiting dilution as well as by flow cytometry. The results showed that the marrows were maintained not as expected by multipotent precursors with enhanced self-renewal, but rather by the collective contribution of lineage-restricted progenitor cells newly endowed with indefinite self-renewal capacity by enforced expression of the homeobox transgene. These findings suggested that the effects of the transduced HOXB4 transgene were manifesting not in early multipotent stages of the hierarchy, but rather most strikingly in later progenitor cells that were terminally committing to individual erythroid or myeloid lineages.

If the immortalizing effect of the HOX transgene was confined to later cells, what then was the means of immortalization of clones initiated by transduced IT-or ST-HSC? Originally infected HSC begin to proliferate and differentiate in culture and subseqently in irradiated mice.

All clonal progeny of infected cells would continue to express the transgene independently of their stage of differentiation. The cells transit initially through early stages of differentiation during which the endogenous Hox genes are expressed alongside the retroviral transgene.

Ultimately, cells in expanding clones reach advanced progenitor stages where the endogenous *Hox* genes are normally silenced. At these stages, progenitors harboring only empty retrovirus would lack capacity for significant in vivo reconstitution. In contrast, in clones initiated by HOX-transduced HSC, HOX protein continues to be available even in late progenitors that no longer express the endogenous *Hox* genes. Our data show that it is these advanced cells that endow the expanding clones with indefinitely extended lifespan in vivo. Our findings are thus consistent with immortalization of clones initiated from transduced IT-or ST-HSC by virtue of the enforced expression of HOX protein in their terminally committed progeny.

In marrows reconstituted from HOX-transduced precursor cells, immortalized cells with the phenotype and lineage-restricted repertoire of late progenitor stages were numerous. Since we used the same infection protocol and transduction vector as used in the original reports of stimulation of HSC expansion by Hox transgenes, we also expected to find elevated numbers of multipotent precursors in reconstituted marrows. However, we were unable to confirm an increase in multipotent precursors of the magnitude described earlier. One possible explanation might be that earlier cells do not respond to exogenous HOX because they already express multiple endogenous *Hox* paralogs. However, because functionally assayed LT-HSC were found not to survive the in vitro transduction protocol, their responsiveness to exogenous HOX expression remains to be formally tested. Nevertheless, it is likely that LT-HSC were present through the initial few days and successfully infected by HoxB4 retrovirus on initial exposure, yet failed to be preserved during further days of culture. The most parsimonious interpretation of the observations is that early HSC stages were not rescued from differentiation or death by the HOXB4 transgene whereas lineage-committed progenitor cells were dramatically affected. The outcome of conditional deletion of the *HOXB4* transgene documented in **Figure 5** provided further evidence for the primary role of late progenitor cells in these recipients. A single transplanted LT-HSC is capable of reconstituting and sustaining blood cell production (Benveniste et al., 2003, 2010). If blood cell production in recipients of *HOX*-transduced cells were sustained by HSC, the grafts would not have been expected to regress on withdrawal of the transgene. The outcome suggests rather that the grafts depended on cells that do not express endogenous *Hox* genes and were thus dependent on exogenous HOXB4 protein for their maintenance. The observations do not support the prevailing view that exogenous HOX proteins have the potential to drive dramatic expansion LT-HSC in culture or in vivo. Based on this view, membrane-permeable HOX proteins have been considered as agents that could enhance the multiplication of LT-HSC numbers in culture prior to clinical stem cell transplantation (Amsellem et al., 2003; Krosl J et al., 2003; Csaszar et al., 2009; Huang et al., 2010). Our findings suggest that the primary action of exogenous HOX protein instead occurs in more advanced progenitor cells in which the endogenous proteins have ceased to be expressed, and should give cause for reconsideration of its rationale in clinical transplantation.

Further information on the stages affected was provided in our study by the failure of transduced HOXB4 to immortalize Progenitor fraction cells. In the two or more days required between exposure to virus and expression of sufficient HOXB4 protein these cells would have differentiated to more advanced stages that may no longer be able to sustain self-renewal even when exogenous HOX protein is made available. Thus it appears that only a narrow window in the differentiation continuum may be susceptible to the immortalizing effect of enforced transgene expression. In agreement, mice reconstituted to 50 - 95% eGFP positivity in all lineages had normal blood counts in all lineages and normal blood cell morphology by conventional microscopy. HoxB4-expressing progenitor cells, and cells with engraftment capacity, remained rare (1 in 1000 - 1 in 15000 marrow cells) within clonally reconstituted mice. Collectively these findings force the conclusion that expression of HOXB4 in amounts sufficient to enable extended self-renewal in progenitor cells does not abrogate differentiation into mature blood cells or interfere with the control points that regulate progenitor frequency or rates of mature cell production.

Enforced expression of an exogenous transcription factor such as HOXB4 could theoretically alter expression of phenotypic markers, and provide an alternative explanation for our failure to detect phenotypic HSC expansion in our experiments. However, the absence of increased numbers of HSC was convincingly supported by the functional analysis which failed to detect transplantable precursors that were fully multilineage in potential. Further arguing against any global interference with marker expression was the detection of the expected committed progenitor phenotypes in elevated numbers, in agreement with their functional detection in transplantation assays. A second theoretical possibility is that exogenous HOXB4 might cause multipotential precursors to appear to be lineage restricted by interfering with choice of particular lineages. However, the committed status of the cells detected in the functional assays was fully supported by our observation of increased numbers of cells with progenitor phenotypes characteristic of lineage-committed precursors. Moreover, such a mechanism is inconsistent with our observations that cells with potential to maintain each of the 4 major lineages were effectively immortalized (**Figures 3 and 4)** ruling out major interference with lineage choice or capacity for appropriate maturation within each lineage. We did find precursors with lymphoid potential to be present at lower frequencies than erythroid and myeloid precursors, consistent with earlier reports of lineage bias within HOXB4-transduced grafts (Kyba et al., 2002; Wang et al., 2005). However, our results favor an explanation on the basis of differing numbers of independently self-renewing myeloid versus lymphoid progenitors, rather than via HOXB4-induced differentiation bias in individual multipotent HSC as has been assumed to now. Further, the fluorescent retroviral expression markers were detected both at progenitor stages as well as in mature differentiated blood cells (Figure 4c and d), indicating that their upstream progenitors in each of the myeloid, erythroid, B and T lineages must also have retained retroviral expression. Thus, differences in abundance between lymphoid and myeloerythroid reconstituting cells appear unlikely to be due to selective silencing of retroviral expression in different lineages.

The coincidence of susceptibility to immortalization with collective cessation of expression of the endogenous Hox cluster genes is illustrated schematically in **Figure 6**. The minimal effect of transgenic HOXB4 on early cell stages that already express endogenous Hox paralogs may reflect sufficiency of the endogenous protein level for maintenance of self-renewal. Published evidence supports the notion that all expressed Hox cluster genes likely contribute to self-renewal of precursor cells. Most Hox cluster paralogs tested by transduction into hemopoietic precursor cells had similar enhancing effects on the ability of cells to reconstitute irradiated hosts. Moreover, the invariant homeodomain common to all the *HOX* genes has been shown to be sufficient for these effects when fused with NUP98 to ensure nuclear localization (Ohta et al., 2007). Our expression analysis newly highlights the coordinated nature of the expression and eventual silencing of multiple HOX cluster genes. Indeed, our data (here and unpublished) together with published RNAseq data (Venkatraman et al., 2013) document active transcription of 15 HOX cluster paralogs (*a2,3,4,5,6,7,9,10*; *b2,3,4,5,6,7,8*; *c6*) in early hematopoietic precursor cells. The observations imply the availability of a pool of diverse HOX paralog proteins contributing to self-renewal of early precursor cells. This extensive functional redundancy likely accounts for the minimal hematopoietic consequences reported after deletion of individual *HOX* genes. Indeed, complete haploid excision of the entire A cluster had surprisingly modest effects (Lebert-Ghali et al., 2010). A large endogenous HOX protein pool may also imply that transduction of any single *Hox* family gene may have only modest impact on the total amount of available HOX protein, whereas transduction into cells that have ceased to express the endogenous genes might be expected to show the most dramatic effect. However, it is possible that individual Hox paralogs may differ from one another in the quality or amount of influence on self-renewal mechanisms, for example by recruitment of differing cofactors (Meis1 for 5’ cluster gene products; PBX1 for 3’ genes including Hoxb4), or in the exact precursor stages that are responsive. It therefor remains to be determined whether our findings with *HOXB4* would be replicated in all details by other Hox transgenes.

**Figure 6.**
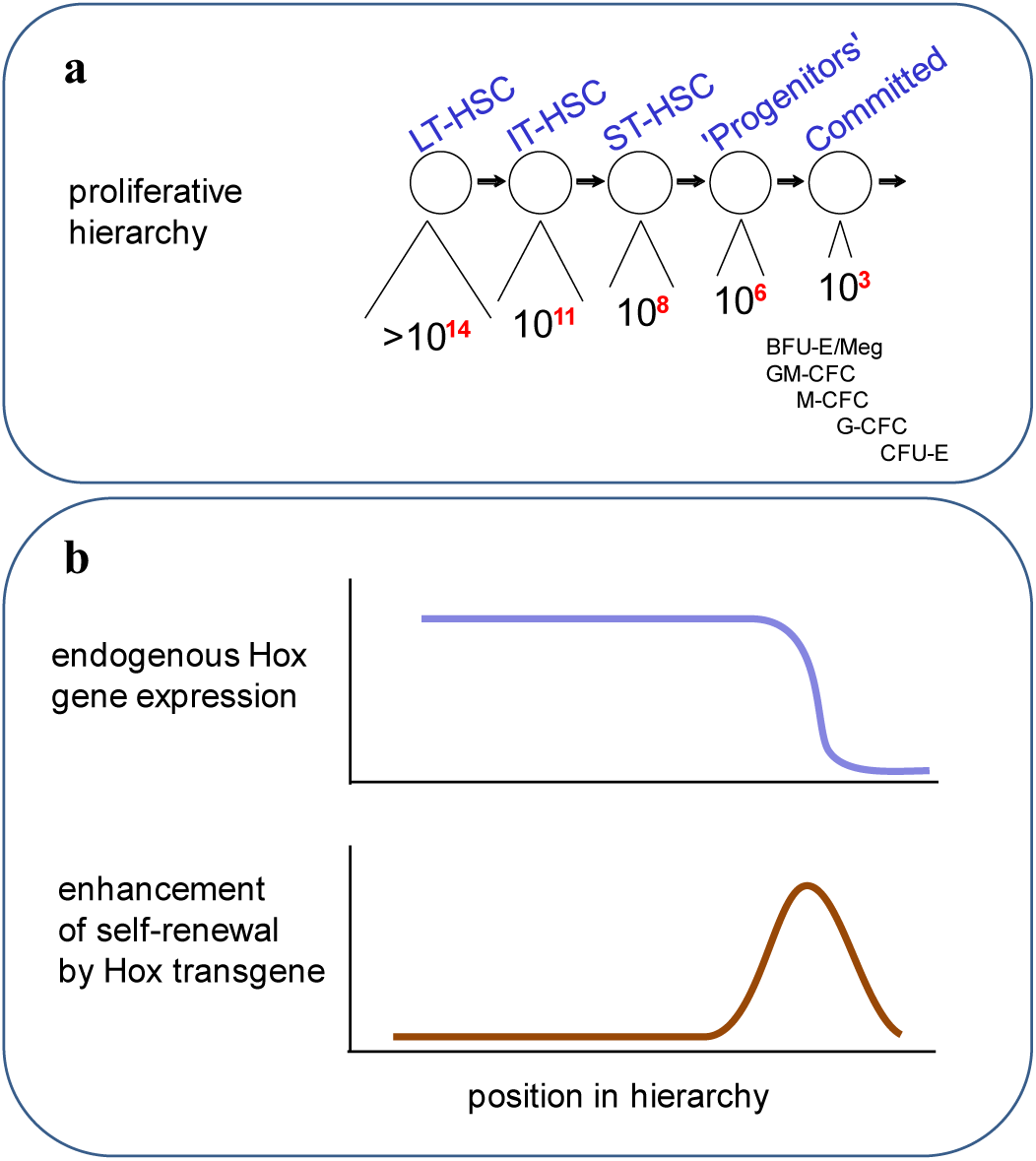
Schematic representation of the relationship in the hemopoietic precursor cell hierarchy between proliferative lifespan, expression of endogenous *Hox* cluster genes and susceptibility to immortalization by enforced expression of an exogenous *Hox* gene. **(a)** Successive stages in the precursor cell hierarchy with estimated maximum attainable clonal size for each stage as established in classical in vivo reconstitution and cytokine-driven in vitro assays. Lineage-committed in vitro colony-forming cells are positioned at the right end of the continuum. **(b)** Aligned against the representation of the hierarchy in (a), susceptibility to immortalization by enforced expression of exogenous HOXB4 coincides with loss of expression of endogenous *Hox* cluster gene expression.

The coincidence in late cells of cessation of endogenous Hox cluster expression and capacity for extension of lifespan by exogenous HOX protein, illustrated in **Figure 6**, implies that it is the shutdown of Hox cluster expression that causes later progenitors to lose the capacity to self-renew. Considered in this light, "self-renewal capacity" in the hemopoietic system appears to be conferred by endogenous HOX proteins in early precursors. The "degree" of self-renewal possible at discrete stages in the hierarchy, and thus the size and longevity of the clones they generate, reflects the amount of time available before the *Hox* clusters are silenced. Once HOX protein is withdrawn, divisions procede without self-renewal until a terminal, post-mitotic stage is reached. The "permanence" of self-renewal possessed uniquely by LT-HSC may reflect special mechanisms that protect the *Hox* genes from shutdown in the LT-HSC lineage within their clones over the course of multiple divisions and extended periods of time. The extensive functional redundancy of the endogenous *Hox* genes has long stymied efforts at conclusive demonstration of their role in early hematopoietic precursor cells. Our core findings bring new and compelling evidence for their centrality in self-renewal of hemopoietic precursor cells.

## MATERIALS AND METHODS

### Mice

Bone marrow donors were C57BL/6J-*Ly5.2-Gpi1*^*b/b*^ or C57BL/6J-*Ly5.1-Gpi1*^*a/a*^ mice. For *in vivo* deletion of retroviral transgene, donor bone marrow cells were taken from B6.129-Gt*(ROSA)26*Sor^tm1(cre/ERT2)Tyj^/J mice. Recipients were ≥8 week C57BL/6J-*Ly5.2-Gpi1*^*b/b*^, C57BL/6J-*Ly5.1-Gpi1*^*a/a*^, C57BL/6J-*Ly5.2-Kit*^*W-41J/W-41J*^*-Gpi1*^*b/b*^, or C57BL/6J-*Ly5.1-Kit*^*W-41J/W-*^*41J-Gpi1*^*a/a*^ mice. Tamoxifen-treated mice were injected intraperitoneally with 100 μL of 10mg/mL tamoxifen (Sigma) dissolved in corn oil for 5 consecutive days. Experiments were conducted in accordance to guidelines and procedures approved by the Animal Care Committee of the Princess Margaret Cancer Centre.

### Purification of Hematopoietic Cells

BM cells were harvested from femurs and tibias of mice, then treated with NH_4_Cl for lysis of red blood cells. For purification of LT-HSC and IT-HSC, NH_4_Cl-treated cells were stained/destained with Rhodamine 123 (Eastman Kodak), lineage depleted (CD5, B220, CD11b, Gr1, 7/4 and Ter-119) using magnetic beads and an AutoMACS separator (MiltenyiBiotec, BergischGladbach, Germany). Lineage depleted cells were incubated with Fc receptor blocking antibody (eBioscience, clone 93), stained with PE-Cy5-Streptavidin or APC-eF780-Streptavidin (eBioscience), PE-Cy7-anti-c-kit (eBioscience, clone 2B8), APC-anti-Sca-1 (eBioscience, clone D7), Pacific-Blue-anti-CD150 (Biolegend clone TC15-12F12.2) and PE-anti-CD49b (BD Pharmingen, clone HMα2) antibodies for 30 min, washed, filtered, and sorted. For purification of ST-HSC and Progenitor fractions, NH_4_Cl-treated cells proceeded directly to lineage depletion by AutoMACS separation without staining/destaining with Rhodamine 123. Lineage depleted cells were blocked with Fc receptor antibody, and then stained with PE-Cy5-Streptavidin or APC-eF780-Streptavidin, PE-Cy7-anti-c-kit, APC-anti-Sca-1, FITC-anti-CD34 (eBioscience, RAM34), and PE-anti-Flt3 (eBioscience, A2F10) antibodies for 30 min, washed, filtered, and sorted.

### Retroviral Vectors & Production of Retrovirus

Non-conditional *GFP-* and human *HOXB4-* expressing MSCV retroviral vectors and production of virus supernatants using GP+E86 producer fibroblasts were as described in (Antonchuk et al., 2001). For experiments involving transgene deletion, the MSCV-related MYs retroviral vector expressing the VENUS fluorescent protein was modified by insertion of loxP sequences within the U3 sequences of the LTR to obtain MYs^loxp^, MYs^loxp^-*HOXB4*, and non-deletable MYs-*Hoxb4*. The human *HOXB4* insert was cloned from the *HOXB4* vector used in (Antonchuk et al., 2001). *Hoxb4* was synthesized from the murine coding sequence and codon optimized for expression. Infectious ecotropic retroviruses were produced by transfection of the vectors into the Plat-E packaging cell line as described in (Morita et al., 2000).

### Cell Culture & Infection of Primary Purified Cells

For clonal studies, purified cells were seeded at an average of 1 cell per Terasaki well in 10 μL IMDM containing 7.5 x 10^−5^ M α-thioglycerol, 4% FBS, 0.1% BSA, 5 μg/ml transferrin, 5 μg/ml insulin, 50 ng/ml c-kit ligand (KL), 50 ng/ml Flt3 ligand (FL), 10 ng/ml interleukin-11 (IL-11), and IL-7 conditioned medium (Benveniste et al., 2003). For ST-HSC and Progenitor cultures, 50ng/mL thrombopoietin and interleukin-3 (IL-3) as 1% conditioned medium from WEHI3B cells were also included. After 16 hr culture to initiate exit from quiescence, entire volumes of wells that were visually confirmed to contain just a single cell were transferred into flat-bottom microtitre wells containing 80-90% confluent irradiated (15 Gy, Cs^137^) GP+E86-GFP or GP+E86-*HOXB4* virus producer fibroblasts (MSCV-IRES-*GFP* and MSCV-*HOXB4*-IRES-GFP vectors (Antonchuk et al., 2001) and 150 μL of the same medium plus cytokines with 4 μg/mL polybrene. After 4 d, loosely adherent and non-adherent cells were recovered from co-cultures and transferred into U-bottom microtitre wells in medium plus cytokines without polybrene for an additional 24 hrs for ST-HSC and Progenitor or 4 days for LT-HSC and IT-HSC. Each well was examined by fluorescence microscopy. Non-adhering cells from clones containing GFP^+^ cells were harvested and passed through a 35 μM mesh filter to remove any non-adhering virus-producer cells before injection into mice. For infection with MYs-based retroviruses, purified cells were cultured overnight before addition of a single dose of retroviral supernatant. Cells were washed after 4 days and entire wells containing VENUS+ cells were transplanted on day 8 into irradiated hosts.

### Reconstitution Assays

For primary transplantations, cultured cells were intravenously injected into the tail vein of sublethally irradiated (4 Gy, Cs^137^) double-congenic C57BL/6J-*Ly5.1-Kit*^*W-41J/W-41J*^*-Gpi1*^*a/a*^ mice. Alternatively, donor cells were co-injected with 1 x 10^6^ unfractionated recipient-type BM cells from C57BL/6J-*Ly5.1-Kit*^*W-41J/W-41J*^*-Gpi1*^*a/a*^ or C57BL/6J-*Ly5.2-Kit*^*W-41J/W-41J*^*-Gpi1*^*b/b*^ mice into lethally irradiated (9 Gy, Cs^137^) C57BL/6J-*Ly5.2-Gpi1*^*b/b*^ or C57BL/6J-*Ly5.1-Gpi1*^*a/a*^ host mice. To evaluate the functional purity of cells following each independent sort, 50 LT-HSCs or 50 IT-HSCs were competed against 1 x 10^6^ unpurified BM cells in lethally irradiated C57BL/6J-*Ly5.2-Gpi1*^*b/b*^ or C57BL/6J-*Ly5.1-Gpi1*^*a/a*^ double-congenic mice. To improve on the sensitivity of detecting ST-HSC and Progenitor cell engraftment, 100s of ST-HSC or 10000s of Progenitors were injected into C57BL/6J-*Ly5.2-Gpi1*^*b/b*^ or C57BL/6J-*Ly5.1-Gpi1*^*a/a*^ mice along with 1 x 10^6^ unfractionated recipient-type BM cells from C57BL/6J-*Ly5.1-Kit*^*W-41J/W-41J*^*-Gpi1*^*a/a*^ or C57BL/6J-*Ly5.2-Kit*^*W-41J/W-41J*^*-Gpi1*^*b/b*^ mice.

For limiting dilution assays, a range of 5 x 10^2^ to 2 x 10^6^ unfractionated BM cells from repopulated primary recipients of Hox-transduced cells were transplanted into a series of lethally irradiated C57BL/6J-*Ly5.2-Gpi1*^*b/b*^ mice along with 1 x 10^6^ BM cells from C57BL/6J-*Ly5.2-KitW-41J/W-41J-Gpi1b/b* mice.

For analysis and secondary transplantation of primitive cells, repopulated BM cells from recipient mice were prepared as described for the purification of LT-HSC and IT-HSC except that cells were not stained with Rhodamine 123, and instead were stained with PerCP-Cy5.5-anti-CD45.1 (eBioscience, clone A20) to mark donor-type cells. CD45.1^+^_GFP^+^_Lin^-^_Sca1^+^_cKit^+^_SLAMF1^+^ or CD45.1^+^_GFP^+^_Lin^-^_Sca1^+^_cKit^+^_SLAMF1^-^ cells were purified, cultured overnight, counted, then transplanted into lethally irradiated C57BL/6J-*Ly5.2-Gpi1*^*b/b*^ mice along with 1 x 10^6^ BM cells from C57BL/6J-*Ly5.2-Kit*^*W-^41^J/W-^41^J*^*-Gpi1*^*b/b*^ mice.

### Detection of Donor Erythroid, Myeloid, and Lymphoid Cells

Peripheral blood was collected from the tail vein or saphenous veins of recipient mice at 2 to 8 week intervals following transplantation. Erythroid reconstitution was measured by quantitation of alternative isoforms of glucose phosphate isomerase (Gpi1^a^ or Gpi1^b^) in blood erythrocyte lysates separated by flat bed electrophoresis (Trevisan and Iscove, 1995). The assay can detect down to 3% of total RBC corresponding to the presence of at least 10^9^ circulating erythrocytes. For tracking of blood leukocytes, erythrocytes were first lysed in NH4Cl blood, blocked with anti-CD16/32 antibody (eBioscience, clone 93), and stained with APC-eF780-anti-CD45.1 (clone A20), PerCP-Cy5.5-anti-CD45.2 (clone 104), PECy7-anti-CD11b (clone M1/70), PECy7-anti-Gr1 (clone RB6-8C5), APC-anti-TCRb (clone H57-597), and eF450-anti-B220 (clone RA3-6B2) or eF450-anti-CD19 (clone 1D3), all from eBioscience. Flow cytometry was performed on a LSRII instrument (BD Bioscience) data were analyzed with FlowJo software (TreeStar Inc.). Donor reconstitution calls in Myeloid, B or T lineages were based on 1) low signal for the other two lineages, 2) contribution of at least 0.05% of the total lineage, and 3) characteristic clustering of events in the same parameter space as donor-only positive controls.

The proportion of donor cells in each lineage was calculated by dividing the number of donor-positive events by the sum of donor-positive and recipient-positive events. BM reconstitution was analyzed with antibodies APC-eF780-Streptavidin, APC-eF780-anti-CD127 (clone A7R34), PE-Cy7-anti-c-kit, APC-anti-Sca-1, Pacific-Blue-anti-CD150, PE-anti-CD49b, eF450-anti-CD34 (clone RAM34), PE-anti-FcγR (clone 93), all from eBioscience.

### Transcript Expression in Purified Cells

cDNA representing various cell populations in the hematopoietic hierarchy was globally amplified as originally described (Brady and Iscove, 1993; Iscove et al., 2002). For LT-and IT-HSCs, cDNAs were generated from 30 - 50 cells immediately following sorting (termed “fresh”) as well as after 48 or 36 hr respectively in culture with KL, FL, IL-11 and IL-7 as above, at which times 50% of cells had divided (Benveniste et al., 2010). For ST-HSC and Progenitor fraction cells, cDNAs were amplified from 30 - 50 cells immediately after sorting as well as after 32 and 24 hr in culture respectively. cDNA from all subsequent stages in the erythromyeloid hierarchy was globally amplified from single cells whose lineage potential was inferred from the fates of sibling cells as described in (Brady et al., 1995; Billia et al., 2001) followed by pooling of biologically similar samples. B and T lymphoid progenitor and mature cell populations (Hardy et al., 2012; Schwarz et al., 2007) were FACS-purified according to the following markers: A (B220+ CD43+ HSA-BP.1-), B (B220+ CD43+ HSA+ BP.1-), C (B220+ CD43+ HSA+ BP.1+), D (B220+ CD43-IgM-IgD-), E (B220+ CD43-IgM+ IgD-), F (B220+ CD43-IgM+ IgD+), DN1 (CD3-CD4-CD8- CD44+ CD25-), DN2 (CD3- CD4- CD8- CD44+ CD25+), DN3 (CD3- CD4- CD8- CD44- CD25+), DN4 (CD3- CD4- CD8- CD44- CD25-), DP (CD4+ CD8+), CD4 SP (CD4+ CD8-), CD8 SP (CD4- CD8+). RNA was extracted from each lymphoid progenitor population by the RNeasy kit (Qiagen), and cDNA was globally amplified. Primer sequences were as follows (5’ to 3’: forward; reverse):

*Hoxb4* (TCATGTGTGTCCTCTCTCCT; CTGTTGTCACTCTGTACAGG), *Hoxa3* (GAGCCTTTTGTTCAATGGTG; TTAGCGTTCAGTTTGGCCAG), *Hoxa4* (TCCCAGCTTTCTAACCTTCC; AAATGCATTTCCCTCTCCCC), *Hoxa5* (CTTGTTCAACGTGTAGTGGC; GCTTAAACAGCCAGACTTGG), *Hoxa9* (GAGCTATACGTGTGTGCAGA; TTTGGTCAGTAGGCCTTGAG), *Hoxa10* (GTCAAACCTGTAGGTGCAGA; TTCCACGCACAGCAGCAATA), *Hoxb6* (TCAATGGTAGATTCGCTGTC; GTATGTGCTCCTTCCAGTGG), *Gapdh* (GCTGGCATTGCTCTCAATGA; AGGCCCCTCCTGTTATTATG), *Hba-a1* (GGACAAATTCCTTGCCTCTG; CAAAGACCAAGAGGTACAGG), *Mpo* (ACCTTCTGGTTGGCAGAAAC; ACAAATAGCACAGGAAGGCC).

All primers were designed to amplify sequences within 300 bp of the true 3’ transcript ends (Muro et al., 2008). Quantitative PCR was performed on the 7900HT Real-Time PCR system (Applied Biosystems) using QuantiTect SYBR Green PCR kit (Qiagen) and analyzed in triplicates. Relative abundance of the target gene was calculated as [(E_Gapdh_)^CtGapdh^] / [[(E_Target_)^CtTarget^], where “E” is the efficiency of amplification of the primer pair over one cycle of PCR, and “Ct” is the number of cycles required to reach a set threshold value of detection of PCR products.

## ACKNOWLEDGEMENTS

We thank Natalie Simard, Gisele Knowles, Pier-Andreé Penttilä and Sherry Zhao for assistance with flow cytometry and Maria-Fernanda Monroy for help with animal procedures. The work was funded by grants from the Terry Fox Foundation, the Canadian Cancer Research Institute, the Canadian Institutes of Health Research and the Stem Cell Network. Additional support came from the McEwen Centre for Regenerative Medicine, the Princess Margaret Hospital Foundation, and the Campbell Family Institute for Cancer Research. This research was funded in part by the Ontario Ministry of Health and Long Term Care. The views expressed do not necessarily reflect those of the OMOHLTC.

